# *Cladocopium* community divergence in two *Acropora* coral hosts across multiple spatial scales

**DOI:** 10.1101/575183

**Authors:** SW Davies, K Moreland, DC Wham, MR Kanke, MV Matz

## Abstract

Many broadly-dispersing corals acquire their algal symbionts (Symbiodiniaceae) ‘horizontally’ from their environment upon recruitment. Horizontal transmission could promote coral fitness across diverse environments provided that corals can associate with divergent algae across their range and that these symbionts exhibit reduced dispersal potential. Here we quantified community divergence of *Cladocopium* algal symbionts in two coral host species (*Acropora hyacinthus, Acropora digitifera*) across two spatial scales (reefs on the same island, and between islands) across the Micronesian archipelago using microsatellites. We find that both hosts associated with two genetically distinct *Cladocopium* lineages (C40, C21), confirming that *Acropora* coral hosts associate with a range of *Cladocopium* symbionts across this region. Both C40 and C21 exhibited extensive clonality. Clones not only existed across host conspecifics living on the same reef, but also spanned host species, reef sites within islands, and even different islands. Both *Cladocopium* lineages exhibited moderate host specialization and divergence across islands. In addition, within every island, algal symbiont communities were significantly clustered by both host species and reef site, highlighting that coral-associated *Cladocopium* communities are structured across small spatial scales and within hosts on the same reef. This is in stark contrast to their coral hosts, which never exhibited significant genetic divergence between reefs on the same island. These results support the view that horizontal transmission could improve local fitness for broadly dispersing *Acropora* coral species.

## Introduction

Many well-known symbioses involve the passing of symbionts from parents to offspring (vertical transmission), fully aligning the evolutionary trajectories of symbiotic partners and typically leading to their deep integration at biochemical and genomic levels (i.e. *Buchnera* in aphids: Nakabachi, Ishida, Hongoh, Ohkuma, & Miyagishima, 2014; Shigenobu & Wilson, 2011). The result of such symbiosis is essentially a novel composite organism, often called the ‘holobiont’, upon which selection can act (Bordenstein & Theis, 2015). In other types of symbioses, the association between partners must be established anew each generation (horizontal transmission), which offers the host’s offspring the opportunity to sample a variety of symbiont lineages and select partners that potentially confer some sort of local advantage (Hilario et al., 2011; Schwarz, Krupp, & Weis, 1999; Usher, Bergman, & Raven, 2007). In theory, this kind of relationship should generate novel ecological opportunities for both symbiotic partners through their mixing and matching across environments. For example, association with ecologically specialized algal photobionts can lead to distinct ecological guilds of lichens (Peksa & Skaloud, 2011) or allow a fungal partner to expand its geographic range across a more broad climatic envelope (Fernandez-Mendoza et al., 2011). Similarly, in aphids, association with various horizontally transmitted bacterial symbionts allows these insects to colonize novel host plants across climatic zones (Henry et al., 2013).

Associations with algal symbionts in the family Symbiodiniaceae are obligatory for the majority of shallow water tropical corals since they rely on photosynthetic byproducts from the algae for energy in oligotrophic waters. In turn, the algae benefit from a protected and light-exposed residence as well as inorganic nutrients and CO_2_ concentration mechanisms provided by the host (Barott, Venn, Perez, Tambutte, & Tresguerres, 2015; Muscatine, 1990; Muscatine & Cernichiari, 1969; Trench & Blank, 1987). Coral symbiosis, like many other ecologically important symbioses, is endosymbiotic (occur within cells) and can establish by two fundamentally different modes of transmission: vertical (symbiont inheritance from mother) and horizontal (symbiont from environmental, free-living sources) (Harrison & Wallace, 1990). Vertically transmitting corals guarantee the maintenance of symbiosis in their offspring, however if larvae encounter novel environments, their symbiont composition may be suboptimal resulting in reduced fitness (Byler, Carmi-Veal, Fine, & Goulet, 2013; Douglas, 1998; Wilkinson & Sherratt, 2001). During horizontal transmission, aposymbiotic larvae have flexibility in symbiont acquisition and upon arrival to new environments, they can uptake novel symbionts not present in parental populations (Abrego, van Oppen, & Willis, 2009; Ali et al., 2019; Gómez-Cabrera, Ortiz, Loh, Ward, & Hoegh-Guldberg, 2008; Little, van Oppen, & Willis, 2004), but availability of symbionts upon arrival is not guaranteed.

Given the obligatory nature of this symbiosis for the host, it is somewhat surprising that in the majority of coral species (∼85%), algal symbionts must be acquired by the juvenile coral from its local environment post settlement (Baird, Guest, & Willis, 2009; Fadlallah, 1983; Harrison & Wallace, 1990; Hartmann, Baird, Knowlton, & Huang, 2017). However, this prevalence of horizontal transmission in coral-algal symbiosis is consistent with a recent meta-analysis on transmission modes in bacteria-eukaryotes being dominated by horizontal transmission in marine environments (Russell, 2019). One possible benefit to this horizontal transmission strategy in marine environments is that these aposymbiotic coral larvae can disperse great distances with ocean currents (Davies, Treml, Kenkel, & Matz, 2015; Foster et al., 2012; Rippe et al., 2017; van Oppen, Peplow, Kininmonth, & Berkelmans, 2011). Yet, coral larvae can encounter a great variety of reef habitats (Gorospe & Karl, 2011), and therefore conditions on the reef where they eventually settle can be very different from their natal reef (Baird, Cumbo, Leggat, & Rodriguez-Lanetty, 2007; LaJeunesse et al., 2004). To improve their chance of survival in this novel environment, corals could potentially associate with locally available, and putatively ecologically specialized, algal strains (Byler et al., 2013; Howells et al., 2012; Rowan & Knowlton, 1995).

Indeed, the diversity of algal symbionts in the family Symbiodiniaceae is rich (LaJeunesse et al., 2018) and specific coral-algae associations have been suggested to play pivotal roles in holobiont adaptation to climate change (Berkelmans & van Oppen, 2006; Howells et al., 2012). The genus *Cladocopium* (formerly clade C *Symbiodinium*; LaJeunesse et al., 2018) originated and diversified most recently among Symbiodiniaceae, and has achieved the highest diversity of all lineages (Lesser, Stat, & Gates, 2013; Pochon & Gates, 2010; Pochon, Montoya-Burgos, Stadelmann, & Pawlowski, 2006; Thornhill, Howells, Wham, Steury, & Santos, 2017; Thornhill, Lewis, Wham, & LaJeunesse, 2014). This diversity has been associated with functional variation in symbiont thermal performance across reefs (Davies, Ries, Marchetti, & Castillo, 2018; Howells et al., 2012) as well as with functional differences in gene expression between reef zones (Barfield, Aglyamova, Bay, & Matz, 2018; Davies et al., 2018), lending support for the potential for reef-specific symbiont communities. In addition, the draft genome of *Cladocopium goreaui* confirm the divergence of this genus from other Symbiodiniaceae genera and specifically highlight that gene families related to the establishment and maintenance of symbiosis (photosynthesis, host-symbiont interactions, nutrient exchange) were under positive selection (Liu et al., 2018).

However, much less is known about the population biology of *Cladocopium* spp. algal symbionts, including how their populations are structured in comparison to their coral hosts. Understanding the relative importance of reef environment, coral host, and geographical distance in structuring coral-associated algal symbiont communities is essential to identifying the adaptive capacity of this symbiosis. However, thus far there are only a handful of population genetic studies of Symbiodiniaceae based on multilocus markers, none of which address all of the above-mentioned potential sources of genetic variation. Here, using microsatellites, we examined the community divergence of *Cladocopium* spp. algal symbionts hosted by two common, co-occurring species of *Acropora* corals– *A. hyacinthus* and *A. digitifera* – collected from the same reef locations across the Micronesian Pacific (Fig 1A, B). We explore this community divergence across several ecological scales including host species, islands across the Micronesian archipelago, and unique sites within each island. We then contrast results for the algal symbionts to the previously published population genetic structure of their coral hosts (based on a subset of the exact same coral samples), which demonstrated that both coral species exhibited extensive genetic connectivity and their genetic structure was well explained by the biophysical connectivity between sites (Davies et al., 2015).

**Fig 1:**
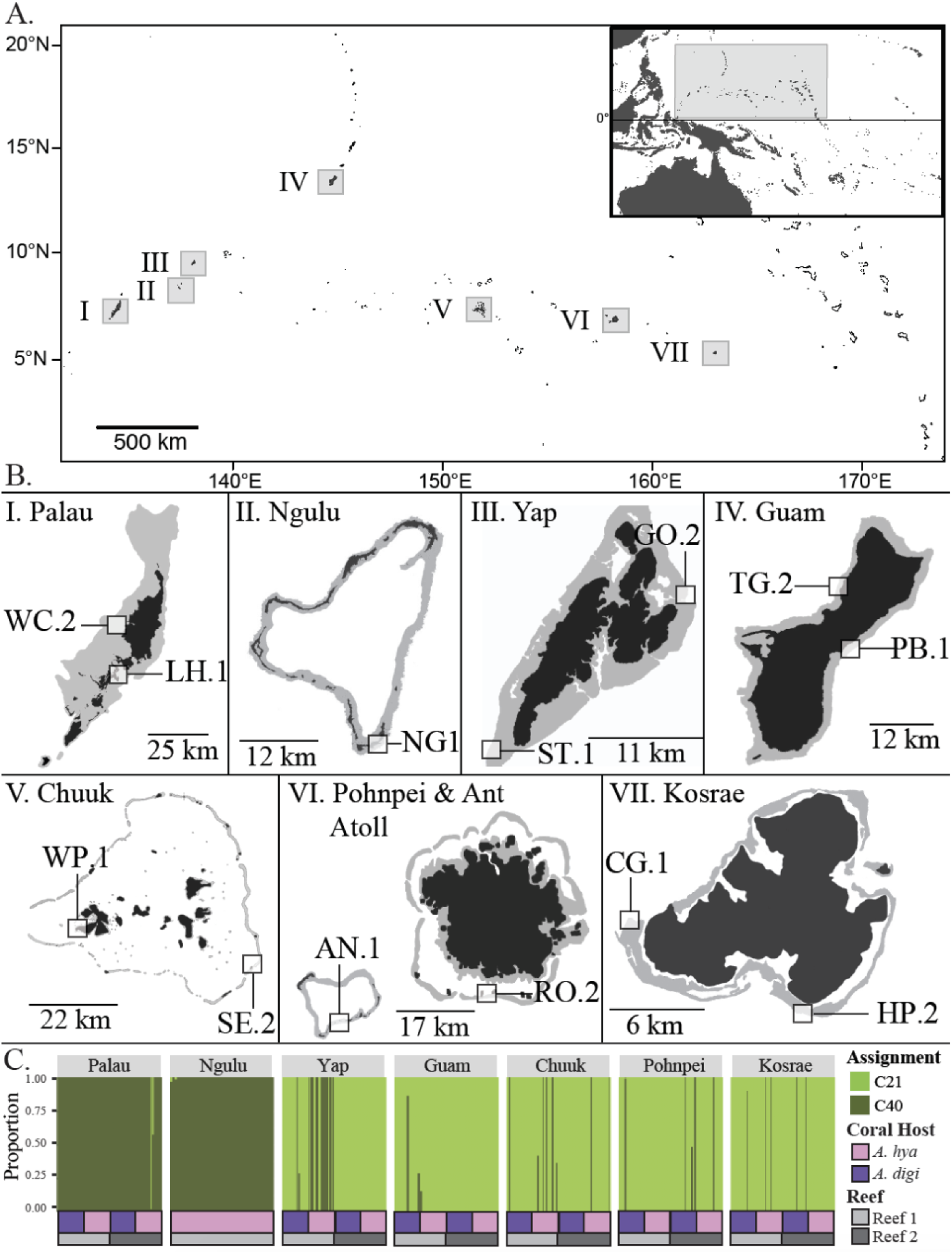
Locations where coral samples were collected and overall DAPC *Cladocopium* community divergence. (A) Sampled islands in Micronesia, with an inset of the Pacific Ocean for reference. (B) Sampled locations within each island. Locations were chosen to potentially maximize within-island divergence. Additional site information can be found in Table 1. (C). DAPC assignments for *Cladocopium* at an optimal cluster number 2, corresponding to C40 (Dark Green) and C21 (light Green). On panel C, color bars below assignment plot indicate coral host species (see legend) and shades of grey correspond to different sites within each island.

## Materials and Methods

### Sampling of coral-associated algal symbionts

This study comprised a subset of samples previously analyzed for coral host genetics in Davies et al. (2015) (Table 1, Fig 1A). Twenty-five individuals of each coral host species (*Acropora hyacinthus* and *Acropora digitifera*) were examined at two reef sites within each of seven islands (Fig 1B). There were two exceptions to this sampling design. First, at Ngulu only one site was visited and only *A. hyacinthus* was collected. Second, at Guam, no *A. hyacinthus* was found on either of the sampled reefs, so only *A. digitifera* was collected. In total, 13 reef sites were included in this experimental design. All samples were collected between 3–7 m depth, all colonies were >2 m apart, and all samples from both species at a given site were collected at the same approximate GPS coordinates (Table 1).

**Table 1:** Reef Site Collections. Site, main island group, GPS coordinates, number of *Acropora digitifera* and *Acropora hyacinthus* hosts genotyped. The first value is the number of individuals successfully genotyped, which were included in the first discrimination analysis (Fig 1C). The second value corresponds to the number of individuals that were successfully discriminated between C3 and C40 at an assignment rate of >0.9 (C40: 172, C3: 388; Fig 1C). Numbers in brackets correspond to the number of individuals hosting unique *Cladocopium* with clones removed, which were included in all downstream analyses (Total: C40: 127, C3: 328; Fig 3, 4). Site letters corresponds to island insets in Figure 1B.

| Site | Island | GPS | *A. digitifera* | *A. hyacinthus* |
| --- | --- | --- | --- | --- |
| WC.2: West Channel | Palau | 7°31'55.7 N, 134°29'42.8 E | 25, 25 (16,0) | 25, 24 (13,1) |
| LH.1: Lighthouse Reef | Palau | 7°16'62.4 N, 134°27'61.9 E | 24, 24 (19,0) | 25, 25 (18,0) |
| NG1: Ngulu | Ngulu Atoll | 8°18'12.0 N, 137°29'18.7 E | 0<sub>1</sub> | 42, 42 (28,0) |
| ST.1: South Tip Reef | Yap | 9°26'05.4 N, 138°02'10.4 E | 25, 24 (1,23) | 25, 25 (0,20) |
| GO.2: Goofnuw Channel | Yap | 9°34'26.4 N, 138°12'19.2 E | 24, 24 (17,5) | 25, 25 (0,24) |
| PB.1: Pago Bay | Guam | 13°25'66.6 N, 144°47'94.3 E | 26, 26 (0,20) | 0* |
| TG.2: Tanguisson | Guam | 13°32'61.1 N, 144°48'52.6 E | 23, 20 (0,17) | 0* |
| WP.1: West Polle | Chuuk | 7°19'69.7 N, 151°33'21.1E | 16, 15 (2,11) | 24, 23 (1,22) |
| SE.2: South East Pass | Chuuk | 7°14'60.3 N, 152°01'29.1 E | 21, 21 (1,20) | 23, 23 (2,13) |
| AN.1: Ant Atoll (East) | Pohnpei | 6°47'42.3 N, 158°01'20.7 E | 24, 24 (0,17) | 24, 23 (3,13) |
| RO.2: Roj | Pohnpei | 6°46'37.7 N, 158°12'24.1 E | 24, 24 (0,21) | 24, 24 (1,21) |
| CG.1: Coral Garden | Kosrae | 5°18'47.2 N, 162°53'01.8 E | 25, 24 (1,19) | 25, 25 (2,20) |
| HP.2: Hiroshi Point | Kosrae | 5°15'88.0 N, 162°59'01.8 E | 25, 25 (2,14) | 25, 25 (0,22) |
| TOTAL |  |  | **282 (73,203)** | **287 (99,185)** |

\* indicates no individuals of this host species were found
<sub>1</sub> indicates individuals were not collected from this site but are likely present

### Laboratory procedures

DNA was isolated following Davies et al. (2013). Microsatellite primers consisted of five previously described *Cladocopium*-specific loci (previously described as clade C *Symbiodinium*) (Bay, Howells, & van Oppen, 2009; Wham, Carmichael, & LaJeunesse, 2014) and one novel locus mined using MsatCommander (Faircloth, 2008) from nucleotide EST data for *Cladocopium* lineage C3 (Leggat, Hoegh-Guldberg, Dove, & Yellowlees, 2007), for a total of six loci (Table S1). Loci were multiplexed in 20 µl polymerase chain reaction (PCR) mixtures containing 10 ng of template DNA, 0.1 µM of each forward and reverse primers, 0.2 mM dNTP, 1X *ExTaq* buffer, 0.025 U *ExTaq* Polymerase (Takara) and 0.0125 U *Pfu* Polymerase (Promega). Amplifications began at 94°C for 5 min, followed by 35 cycles of 94°C for 40 s, annealing temperature for 120 s, and 72°C for 60 s and a 10 minute 72°C extension period. Molecular weights were analyzed using the ABI 3130XL capillary sequencer. Data were binned by repeat size and individuals failing to amplify at ≥3 loci were excluded from downstream analyses.

### Analyses of allele presence-absence data

Although Symbiodiniaceae *in hospite* are assumed to be haploid (Santos & Coffroth, 2003), the genus *Cladocopium* are generally observed to have two copies of every allele (Thornhill et al., 2014; Wham et al., 2014; Wham & LaJeunesse, 2016). This apparent genome duplication may or may not correspond to a change in chromosome number, or the actual diploid state (Wham & LaJeunesse, 2016), and it has been previously suggested that these lineages should be scored as if they were effectively diploid (i.e. with the expectation of two alleles per locus) to appropriately construct multilocus genotypes (MLGs) from samples (LaJeunesse et al., 2014; Pettay, Wham, Smith, Iglesias-Prieto, & LaJeunesse, 2015; Thornhill et al., 2014; Wham et al., 2014; Wham & LaJeunesse, 2016). However, ploidy of the *Cladocopium* samples in our study is unknown, and a single coral could potentially contain several genetically distinct *Cladocopium* clones. Therefore, data were analyzed as binary (allele presence/absence) values for each sample. This conservative approach allowed us to retain all individuals in analyses (569 total: 282 *A. digitifera* and 287 *A. hyacinthus*), with the drawback that it confounds genetic divergence and community divergence in cases of multiple strains per host. However, since multiple-strain infections are rare in *Cladocopium* (Thornhill et al., 2017), genetic divergence is expected to be the major contributor to our distance measures. Still, we chose to refer to our distances as “*Cladocopium* community divergence” throughout this manuscript, to ensure that there is no confusion with true genetic distances.

First, all binary allele data (N=569 samples: Supplemental File S1) were converted into a *genind* object with allele presence/absence data using *adegenet* 2.0.0 (Jombart, Devillard, & Balloux, 2010) in R (R Development Core Team, 2018). Next, discriminant analysis of principal components (DAPC) was implemented, which classifies samples into user-defined groups based on their coordinates in principle components’ space. Because DAPC does not rely on traditional population genetics models, it is free from Hardy-Weinberg equilibrium and linkage disequilibrium assumptions and thus is considered to be applicable across organisms regardless of their ploidy and genetic recombination rates (Jombart et al., 2010). Here, identification of clusters was achieved using the *find.clusters* function with a maximum number of 40 clusters. Eighteen principle components (PCs) were maintained and the Bayesian Information Criterion (BIC) indicated that two clusters were optimal in our data. In this initial analysis of all data, samples exhibited strong assignments into two highly supported clusters - Light green and Dark green (Fig 1C). These data were therefore split into two subsets, corresponding to these two clusters, for downstream analyses. Only samples assigned to one of the two clusters with a probability >0.9 were retained, resulting in N=388 for the Light green cluster and N=172 for the Dark green cluster (Table 1, Supplemental Files S2 and S3).

### *Sequencing analysis of* Cladocopium *psbA*_*ncr*_

To confirm phylogenetic affiliation of the two highly-supported *Cladocopium* clusters, the non-coding region of the psbA chloroplast gene (psbA_ncr_) was amplified in representative samples. The psbA_ncr_ region was chosen because of its utility for differentiating species of Symbiodiniaceae (i.e. Lewis, Chan, & LaJeunesse, 2019). Amplifications were conducted using the primers and settings described by Moore et al. (2003). Amplified products were directly sequenced using the reverse primer. Phylogenetic analysis was conducted using PAUP Version 4.4a147 (Swofford, 2014) using psbA_ncr_ reference sequences for C40 (from various scleractinians) and C21 (from Acropora) provided by the LaJeunesse lab. The nexus file used to generate the phylogenetic tree containing these reference sequences is included as Supplemental File S4. An unrooted distance tree was constructed to demonstrate that the representative samples from the two highly-supported clusters are separated by large differences in sequence divergence. These results indicated that the two major clusters by our genetic data were C40 (*sensu* LaJeunesse et al., 2004) and C21 (*sensu* Thornhill et al., 2014), respectively. Further community divergence analysis of these data were completed for each cluster separately and these lineages are referred to as *Cladocopium* C40 and *Cladocopium* C21 throughout the rest of the paper.

### Clonal analyses within each cluster

To determine rates of clonality within *Cladocopium* C40 and C21, individuals with matching MLGs were investigated. Clonal analyses for C40 and C21 were conducted independently using the following steps. Singleton alleles were removed (12 alleles in C40; 10 alleles in C21), after which a hierarchical clustering tree was constructed in R (R Development Core Team, 2018) using the *vegdist(x, binary=T, method=“manhattan”)* function from the *vegan* package (Oksanen et al., 2013) and processed with the function *hclust(x)*. Manhattan distance was chosen for this analysis because it corresponds to the total number of non-shared alleles between two MLGs, which should be zero for identical MLGs (clones). Samples sharing identical MLGs were then identified using the function *cutree(x, h=0.2)*, which grouped individuals with less than one (i.e., zero) non-shared alleles. The probability of chance occurrence of matching MLGs was assessed by a resampling simulation in R. 100,000 artificial MLGs were generated using 44 non-singleton alleles in C40 and 46 non-singleton alleles in C21, by resampling the vector of presences-absences of each allele across individuals. The probabilities of identical genotype groups occurring by chance were calculated using the formula 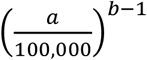, where *a* is how often the MLG was observed in the simulation and *b* is how often it was observed in the actual data. For downstream DAPC analyses, we have created dataframes including only a single representative of each MLG within a site within the same host species (Supplemental Files S5 and S6 for C40 and C21 respectively).

The geographical distance spanned by MLGs was investigated by calculating a distance matrix from reef site coordinates, in decimal degrees, using the *dist* function in R. The *dms2dec* function from Zanolla et al. (2018) was used to convert degrees minutes seconds to decimal degree format. The largest distance was taken for MLGs spanning more than one site. Distances were converted to kilometers using the National Hurricane Center and Central Pacific Hurricane Center’s calculator (https://www.nhc.noaa.gov/gccalc.shtml). Genotype diversity, the probability that two randomly sampled MLGs were different, was calculated using the formula 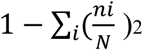, where *n*_*i*_ is the number of individuals representing a *i*’th repeated MLG and *N* is the total number of individuals. Genotypic diversity differences between C40 and C21, and between host species within C40 and C21 were tested using *t.test()* in R.

### Within-cluster analyses across coral hosts, islands and sites within islands

To visualize *Cladocopium* community divergence between host species, between islands, and between sites and host species within each island, assignment of samples to genetic clusters with prior grouping of island/host/site was performed in R (R Development Core Team, 2018) using DAPC (Jombart, 2008; Jombart et al., 2010) separately for C40 and C21. Successful reassignment, indicated by a high proportion of samples correctly assigning back to their *a priori* group, indicates that these user-defined groups are distinct, which in our case implies divergence between hosted *Cladocopium* communities. Here, data were converted into principal components (PCs) and then a-scores were computed to determine the optimal number of PCs to retain. A-scores determine the proportion of successful reassignment corrected for the number of PCs retained and protect against model overfitting (Jombart et al., 2010). Assignment rates, PCs and DFs retained, and the overall proportion of variance explained by each of the models are included in Table S2. Proportion of successful assignments within each model are shown on all figures (Fig 3, 4).

**Fig 2:**
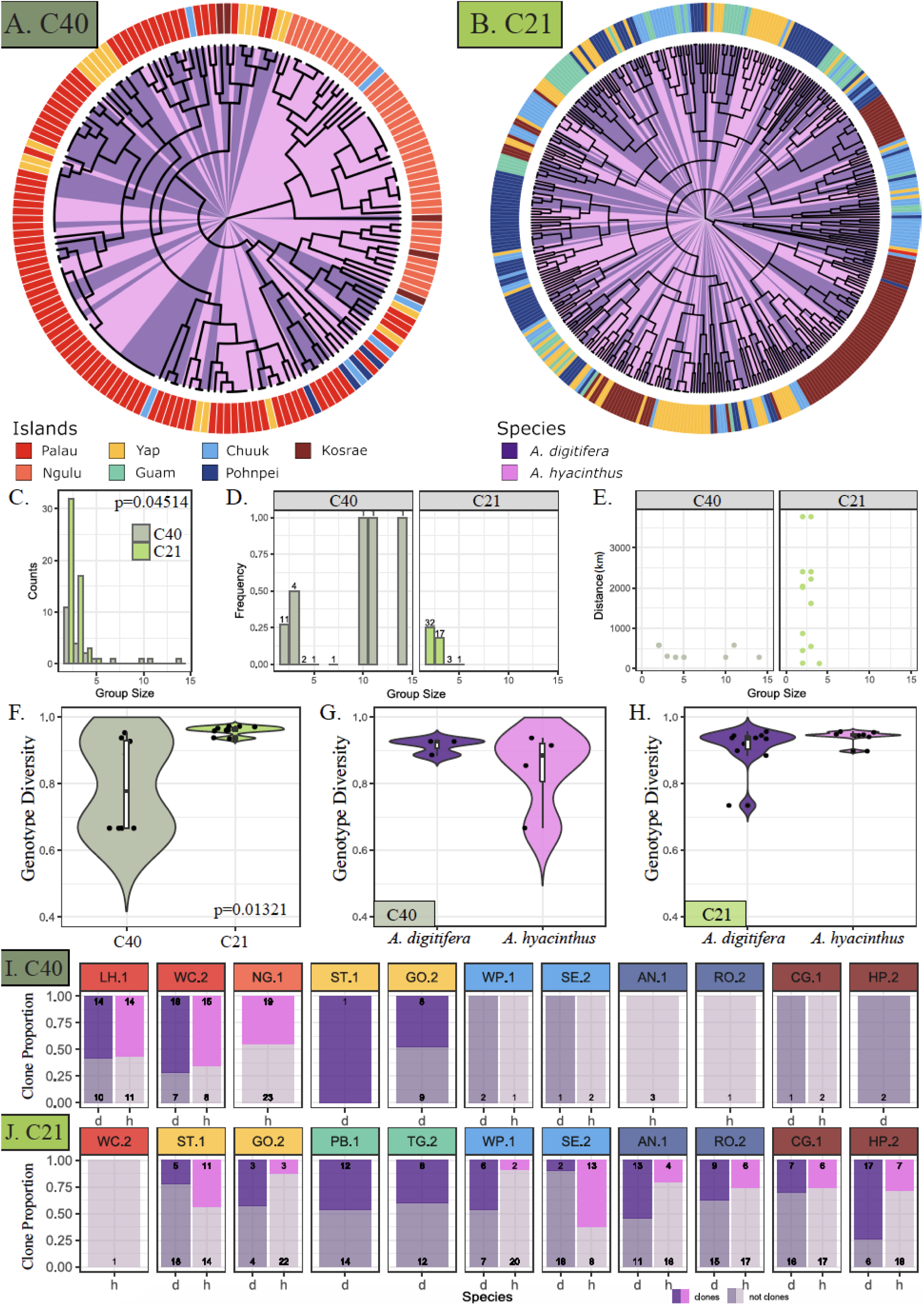
Rates of *Cladocopium* clonality within C40 and C21. Fan trees of *Cladocopium* C40 (A) and *Cladocopium* C21 (B) MLGs. Host species, *A. hyacinthus* and *A. digitifera*, are color-coded on the inside of the tree and the seven islands in Micronesia are indicated in the ring around the tree. (C) Histogram of MLG group sizes. C40 tends to have larger MLG groups than C21. (D) Frequencies of MLGs spanning host species, binned by MLG group size. The number indicates the total number of MLG groups in the size bin. There is no clear difference in the proportion of host-spanning MLGs between C40 and C21. (E) Greatest geographical distance spanned by a MLG group of a given size. C21 MLGs span considerably larger distances than C40 MLGs. (F-H) Violin plots of genotype diversity (probability that the two randomly picked MLGs are different) for C40 and C21 (F) and for each cluster among two host species (G, H). Each point represents a reef site. (I-J) Clonality levels at each reef site in each host species. Bar colors correspond to host species, where faded bar segments represent unique MLGs and bright bar segments represent repeated MLGs. h, *A. hyacinthus*; d, *A. digitifera*. Reef site colors correspond to Fig 1 and Table 1.

**Fig 3:**
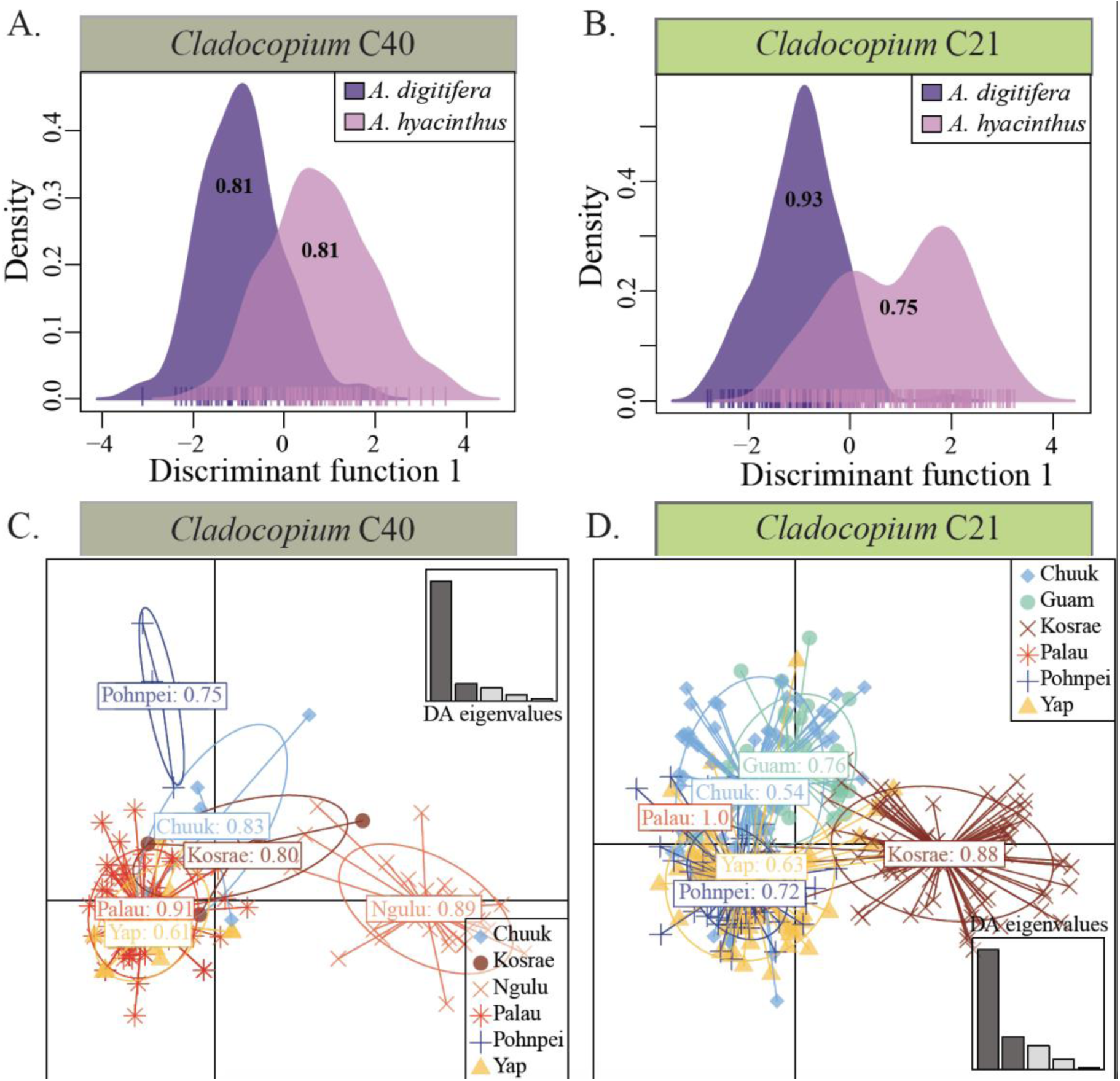
DAPC of binary MLG data for *Cladocopium* C40 and C21 by host species and islands. Discriminant analysis of principal components (DAPC) of binary MLG data for *Cladocopium* C40 and C21 hosted by *Acropora hyacinthus* and *A. digitifera* at thirteen sites across seven islands in Micronesia. DAPC analysis on two discriminant functions demonstrating strong host species assignments across all islands for C40 (A) and C21 (B). Numbers overlaying the curves indicate proportions of correctly assigned samples. DAPC scatter plot for individual samples from C40 (C) and C21 (D) represented by colored dots clustered by islands. Proportions of correct assignments are indicated within the clusters. Information on the DAPC models can be found in Table S2.

**Fig 4:**
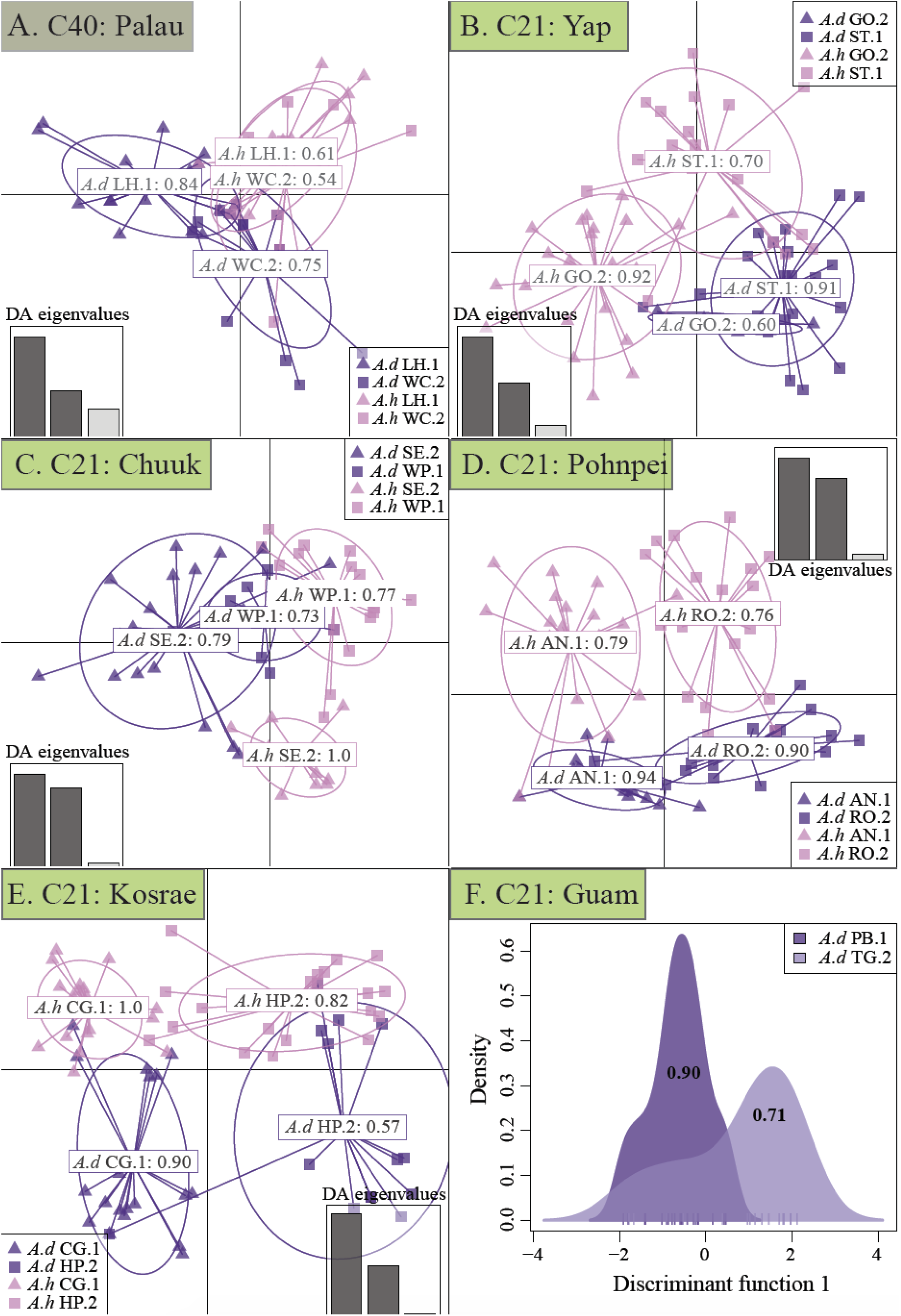
DAPC of *Cladocopium* C40 and C21 hosted by *Acropora hyacinthus* and *Acropora digitifera* at twelve reef sites across six islands in Micronesia. Discriminant analysis of principal components (DAPC) of binary MLG data for *Cladocopium* C40 and C21 hosted by *A. hyacinthus* and *A. digitifera* at two sites within each island in Micronesia. The first two discriminant functions are shown, which generally correspond to host species and site assignments. DAPC scatter plots for individual samples from within (A) Palau for C40, (B) Yap for C21, (C) Chuuk for C21, (D). Pohnpei for C21, and (E) Kosrae for C21. Density plots are shown for the two sites in Guam for C21 for *A. digitifera* hosts only (purple distributions) (F). Proportions of correct assignments are indicated in the clusters and information on the DAPC models can be found in Table S2.

### *Unconstrained* Cladocopium *community analyses*

Because DAPC analyses aim to maximize variation between pre-defined groups, we also visualized all C40 and C21 data independently using a principal coordinate analysis of allele presence/absence data using the *vegdist (x, method=“bray”)* function implemented in the *vegan* package in R (Oksanen et al., 2013).*Cladocopium* community divergence between host species, islands and host species and sites within islands were then tested using the *vegan::adonis* function (*method=“bray”*).

### Data and code availability

All data and code used for all analyses and figure generation are publicly available at https://github.com/daviessw/Cladocopium_Micronesia.

## Results

### *Two clusters of* Cladocopium *symbionts observed in Micronesian acroporids*

Across the two coral host species in Micronesia (Fig 1A, B), two distinct *Cladocopium* clusters were observed with 98.4% of samples (560/569) strongly assigning to one of the two clusters (Fig 1C). Sequencing of the psbA_ncr_ gene from representative samples from each cluster identified them as *Cladocopium* C40 and C21 (LaJeunesse et al., 2004; LaJeunesse & Thornhill, 2011; Thornhill et al., 2014) (Fig S1). It is important to note that the possible presence of other background Symbiodiniaceae genera would not affect *Cladocopium* genotyping results since our microsatellite assays are genus-specific (Bay, Howells, & van Oppen, 2009; Wham, Carmichael, & LaJeunesse, 2014). Corals of both *Acropora* species from Palau and Ngulu were found to almost exclusively host *Cladocopium* C40 (Fig 1C, dark green bars). C40 was also prevalent in *A. digitifera* at one reef site on Yap (Goofnuw Channel: GO.2) and was occasionally found in *A. digitifera* across the rest of Micronesia (Fig 1C). All other *Acropora* hosts across the spatial gradient associated with *Cladocopium* C21 (Fig 1C, light green bars). Both *Cladocopium* lineages possessed high allelic diversity, with a total of 44 unique alleles in C40 (N=127 corals) and 49 unique alleles in C21 (N=328 corals).

### *Clonality of* Cladocopium *symbionts*

Overall, clonality was high in both C40 (51.7%; N=127) and C21 (34.0%; N=328), however rates of clonality in C40 were significantly higher than C21 even though more corals hosted C21 across the archipelago (*p*=0.04514; Fig 2A-C). C40 was represented by a total of 105 unique MLGs, 22 of which were found in multiple individuals (Fig 2A). In C21 there were 309 unique MLGs, 53 of which occurred repeatedly (Fig 2B). 16/22 C40 MLGs and 53/53 C21 repeatedly occurring MLGs were supported with high confidence (*p*<0.001) as clonal groups via simulation. Six repeated MLGs in C40 were less robustly supported (group 12: *p*=0.0166, N=2; group 14: *p*=0.0044, N=2; group 32: *p*=0.0014, N=2; group 35: *p*=0.0173, N=2; group 81: *p*=0.0395, N=2; group 87: *p*=0.0017, N=2), but still all passed the *p*<0.05 significance threshold.

Notably, repeated MLGs were not restricted to a specific host species or reef site. Many spanned host species, reef sites, and even islands (Fig 2A,B). Summaries of clonal rates for each reef site are shown in Fig 2 I-J. MLG group size was on average 4.05 for C40 and 2.49 for C21, ranging up to 14 in C40 and 5 in C21 (Fig 2C,D). Consistent with larger MLG group sizes in C40 relative to C21 (*p*=0.04514; Fig 2C), C40 also exhibited lower overall genotype diversity than C21 (*p*=0.01321; Fig 2F). Notably, larger MLG group size in C40 did not translate into larger geographic distance spanned by an MLG (Fig. 2E). The largest distance spanned by C40 MLGs was between Goofnuw Channel (GO.2), Yap and Lighthouse Reef (LH.1), Palau (∼578 km), while the largest distance spanned by C21 MLGs was between South Tip (ST.1), Yap and Hiroshi Point (HP.2), Kosrae (∼3714 km) (Fig 2E). Rates of MLGs spanning host species also varied between C40 and C21. 36.4% of MLGs spanned host species in C40 (Fig 2A) and 20.8% of MLGs spanned host species in C21 (Fig 2B). There was no significant difference in genotype diversity between algal symbionts hosted by different host species for either C40 (*p*=0.3403; Fig 2G) or C21 (*p*=0.1619; Fig 2H).

### Cladocopium *community divergence by coral host species, islands, and local reef environments*

Discriminant analysis of principal components (DAPC) strongly differentiated between host species for both *Cladocopium* C40 and C21 (Table S2, Fig 3A,B), with assignment rates ranging from 0.75 (*A. hyacinthus* hosting C21) to 0.93 (*A. digitifera* hosting C40). In addition, unconstrained analyses confirmed these results with significant differences observed in both C40 (*adonis* p<0.001) and C21 (*adonis* p<0.001) (Fig S2A,B). These results confirm that host species play a role in structuring *Cladocopium* populations across Micronesia. However, both coral hosts were capable of maintaining symbiosis with multiple Symbiodiniaceae lineages (C40 or C21) across this region. In addition, DAPC demonstrated clustering among islands for each *Cladocopium* species irrespective of host species: generally high per-island assignment rates were obtained both for C40 (Fig 3C, 0.61-0.91) and C21 (Fig 3D, 0.53-0.88), which were also confirmed with significant differences for both C40 (*adonis* p<0.001) and C21 (*adonis* p<0.001) across islands using unconstrained analyses (Fig S2C,D). Notably, algal symbionts from Yap consistently showed some of the lowest assignment rates for both C40 (0.61) and C21 (0.63). Another notable observation was that algal symbionts from Ngulu and Kosrae were strongly separated from other algal communities, suggesting the possibility of additional *Cladocopium* lineages existing there (Fig 3C,D), which was not further explored here.

When clustering was performed within islands for C40 (Palau) and C21 (Yap, Chuuk, Pohnpei, Kosrae, Guam), of the two top eigenvalues in DAPC analysis, generally one discriminant function (DF) explained divergence by host species while the other DF explained differences between reef sites (Fig 4). Unconstrained analyses corroborated this result: *Cladocopium* communities were always significantly different between coral host species and sites within islands (Fig S3). One instance where DAPC clustering and unconstrained analysis by sites and host species within island showed weaker support was C21 from Yap (Figs 4B, S3B), however, the sample size of *A. digitifera* symbionts at Goofnuw (GO.2) was low (N=5). This low sample size was due to *A. digitifera* hosts at Goofnuw largely associated with C40 (N=17), while the majority of other *A. digitifera* hosts across Yap associated with C21 (Fig 1C), which may have led to lack of power to detect clustering. The strongest separation between host:site groups were observed at Chuuk (Figs 4D, S3D) and at Kosrae (Figs 4E, S3E).

## Discussion

### Acropora *corals establish symbiosis with distinct* Cladocopium *communities*

Across the Micronesian Pacific (Fig 1A), both *Acropora* coral hosts associated with two distinct lineages of *Cladocopium* (Fig 1C), which were identified as C40 and C21 (Fig S1), with potentially additional species present (i.e. Ngulu *Cladocopium* C21; Fig 3C). This observation suggests that both coral hosts show considerable flexibility in their symbiotic associations with *Cladocopium* across their range and within their specific environments (Abrego et al., 2009; Berkelmans & van Oppen, 2006). This association with *Cladocopium* is consistent with the wealth of previous comprehensive community composition studies suggesting that Indo-Pacific acroporids are generally dominated by algal symbionts in this genus (i.e. LaJeunesse et al., 2004; LaJeunesse et al., 2003; Thornhill et al., 2014). While initial symbiont infection is likely determined by local availability of free-living symbionts and made possible by the flexibility of arriving coral recruits (Abrego et al., 2009; Ali et al., 2019; Cumbo, Baird, & van Oppen, 2013; Little et al., 2004), a winnowing process involving selection, competition, immune system response, and differing rates of asexual proliferation likely play roles in the eventual dominance of a unique algal community in the majority of coral hosts in their respective habitat (Rowan, Knowlton, Baker, & Jara, 1997; Thornhill et al., 2017).

Strict associations with a single Symbiodiniaceae algal lineage have been observed across a variety of coral hosts and Symbiodiniaceae genera (Baums, Devlin-Durante, & LaJeunesse, 2014; Pinzón, Devlin-Durante, Weber, Baums, & LaJeunesse, 2011; Thornhill et al., 2014), however this strict association is not always the case (see Howells, van Oppen, & Bay, 2009; Howells, Willis, Bay, & van Oppen, 2013). In our study, it is also important to acknowledge that we only explored community divergence patterns within *Cladocopium* because we leveraged *Cladocopium-*specific microsatellite loci (Bay, Howells, & van Oppen, 2009; Wham, Carmichael, & LaJeunesse, 2014). This genus is most commonly known to associate with *Acropora* in this region, which is consistent with our previous ITS2 metabarcoding results on the same coral samples from Palaun reefs, which showed that *Acropora* hosts strictly associated with one of two *Cladocopium* symbiont haplotypes (Quigley et al., 2014). Here, we tested several samples (N=4) for community level algal species identification (Fig S1), which confirmed C40 and C21 designations, however these more coarse-grained genus-level analyses were not performed on samples from across the range. Therefore, we are unable to comment on other algal genera known to inhabit corals at background levels (Silverstein, Correa, & Baker, 2012; Ziegler, Stone, Colman, Takacs-Vesbach, & Shepherd, 2018).

### Cladocopium *C40 and C21 exhibit extensive clonality*

Research leveraging microsatellite loci in Symbiodiniaceae demonstrate that rates of clonality can substantially differ between Symbiodiniaceae genera, between lineages within a genus, and between regions (Thornhill et al., 2017). For example, work on *D. trenchii* hosted by *Acropora* colonies found very low rates of shared clones between colonies (Hoadley et al., 2019), and similarly low clonality rates have been observed in *Dusurdinium* from *Galaxea fascicularis* from the South China Sea (Chen et al., 2020). However, Pettay et al. (2011) found that unique *Pocillopora* hosts frequently associated with the same *S. glynni* clone. Here we find high frequencies of clonality in *Cladocopium* symbionts in *Acropora* hosts with 51.7% of corals hosting C40 sharing the same MLG (Fig 2A) and 34% in C3 (Fig 2B), which cannot be explained by clonality of coral hosts (Davies et al., 2015). While few studies have investigated clonality in Pacific *Cladocopium*, Howells et al. (2013) found that only 13% of *A. millepora* from the Great Barrier Reef hosted the same MLGs. However, clonality rates appear to be different across *Cladocopium* lineages. For example, Thornhill et al. (2014) observed that C3 hosted by *Siderastrea siderea* exhibited 17% clonality, C7 hosted by *Orbicella* spp. exhibited 70% clonality, and C7a/C12 hosted by *Orbicella* spp. exhibited 47% clonality. In light of these data, our observed clonality rates for C40 and C21 are well within previously published estimates.

Interestingly, we found that *Cladocopium* clones were not only shared across conspecifics on the same reef, but also across different host species, different reefs on the same island, and even between host species on different islands (Fig 2A,B). Given that unique MLG genotypes have been shown to exhibit functional variation both in culture (i.e. *S. psygmophilum*, Parkinson et al., 2016) and in hospite (Davies et al., 2018; Howells et al., 2012), these results are counter intuitive for several reasons. First, it is difficult to imagine how clones are distributed across such wide spatial scales, which was especially evident in C21 (Fig 2E), given that the majority of symbioses in corals are horizontal (Baird et al., 2009) and Symbiodiniaceae are expected to have low dispersal potential (reviewed in Thornhill et al., 2017). Secondly, it seems unlikely the same C40 or C21 clone would be successful across both host species and across different environments given that coral-associated symbiont distributions have been proposed to correlate with depth (Andras, Kirk, & Harvell, 2011; Kirk, Andras, Harvell, Santos, & Coffroth, 2009), temperature (Baums et al., 2014; Hume et al., 2016; LaJeunesse et al., 2014), PAR (Rowan et al., 1997), and host species (Thornhill et al., 2017; Thornhill et al., 2014). Another interesting discussion point is that C21 clones appear to be more broadly distributed across the seascape than C40 clones, suggesting that C21 may have higher dispersal potential than C40. Clearly, more intensive sampling of hosts associated with *Cladocopium* across additional host species and sites will shed light on the population biology of these generalist symbionts.

### Cladocopium *C40 and C21 exhibit imperfect host specificity*

The majority of reef-building coral species associate with a specific Symbiodiniaceae type, which have traditionally been coarsely defined based on ribosomal and/or chloroplast markers (Fabina, Putnam, Franklin, Stat, & Gates, 2013; Rodriguez-Lanetty, Krupp, & Weis, 2004; Thornhill et al., 2014; Weis, Reynolds, deBoer, & Krupp, 2001). Previous Symbiodiniaceae multilocus genotyping studies revealed that each of these symbiont types can harbor within-type diversity, both at genetic and functional levels (Howells et al., 2012; Howells et al., 2009; Santos, Shearer, Hannes, & Coffroth, 2004). Here we observe significant divergence between *Cladocopium* communities among two different host species in both C40 and C21 across the Micronesian Pacific (Fig 3A,B; Fig S2A,B), and this pattern of host specificity consistently holds between host species on the same reef (Fig 4; Fig S3). Previous work on octocorals similarly observed significant host differentiation among algal symbionts, however they found that this genetic divergence was driven by different aged cohorts and depth in their system (Andras et al., 2011). Here, host habitat depth or age class is not relevant for the host specificity observed given that specific attention was paid to collecting colonies located at similar depths and of similar size classes. Instead, our data suggest that for both C40 and C21, local association of hosts and symbionts within the same cluster is due to host specificity in *Cladocopium* (Fig 4; Fig S3), which has been previously proposed in symbionts hosted by *Pocillopora* in the south Pacific (Magalon, Baudry, Husté, Adjeroud, & Veuille, 2006). Since our study rigorously sampled two coral host species across several spatial scales, we also detected that this specificity is imperfect: at every location, there were symbionts in one host species that would have been assigned to another coral host based on their MLG (Fig 4, Fig S3). In fact, there were MLGs both within C40 and C21 that were shared across hosts at the same site and across different islands (Fig 2A,B), further highlighting that this host specificity is imperfect. Overall, these patterns suggest that host specialization in *Cladocopium* is present, however the boundary between hosts appears permeable in *A. hyacinthus* and *A. digitifera* across the spatial scale investigated here.

### *Divergent* Cladocopium *communities within islands*

Within each island and sympatric host species, all *Cladocopium* pairwise comparisons exhibit high assignment rates back to their *a priori* groups (Fig 4), which demonstrates significant community divergence between closely located reef sites (Fig S3). It is tempting to speculate that *Cladocopium* community divergence among individual reefs might be due not only to dispersal limitation, but also to spatially varying selection, implying environmental specialization (i.e. local adaptation) in the symbionts. Unfortunately, these islands are remote and understudied and therefore we cannot provide further support for this claim as we did not measure environmental parameters and did not assess symbionts’ fitness across environments. Among factors that might contribute to genetic subdivision among reefs irrespective of distance is high variation in reproductive success among *Cladocopium* clones on a local scale, which would elevate divergence due to spatial discordance of short-term allele frequency fluctuations (Thornhill et al., 2017). Yet, previous work has demonstrated that other *Cladocopium* symbiont populations have exhibited classic signals of local adaptation (Howells et al., 2012), and therefore reef sites investigated here offer an excellent study system for investigating the fine-scale local adaptation potential of *Cladocopium*. If these algal symbionts are indeed locally adapted, this would ensure that horizontally transmitting coral hosts increase their local fitness by associating with local symbionts. To confirm this hypothesis, future work is required to experimentally demonstrate that these symbionts are achieving their maximum fitness in their local reef environment (Kawecki & Ebert, 2004).

### Cladocopium *communities are more spatially structured than their coral hosts*

With our conservative approach to analysis of our symbiont genetic data we cannot directly compare the divergence of symbiont communities to the previously published genetic structure of their coral hosts (Davies et al., 2015). Still, we can compare these results qualitatively. For symbiont communities hosted by the same coral species, we consistently find significant divergence between different sites within the same island (Fig 4; Fig S3). In contrast, no significant within-island genetic divergence was ever detected for either host species, using the exact same coral samples (Davies et al., 2015). This indicates that C40 and C21 algal symbiont communities are more spatially structured that their coral hosts across the same spatial scale.

Strong community divergence in *Cladocopium* was not surprising given the prevailing view of their life cycle. It involves symbiotic existence in sedentary hosts alternating with a short-term free-living form that largely exists in the benthos. The opportunity for *Cladocopium* dispersal by ocean currents is therefore limited, and the primary role of the free-living stage is to invade novel hosts (Fitt, Chang, & Trench, 1981; Fitt & Trench, 1983; Littman, van Oppen, & Willis, 2008; Magalon et al., 2006; Yacobovitch, Benayahu, & Weis, 2004). Our data support this hypothesis with the observation of significant clustering between all pairs of sampled sites within islands in both C40 and C21 lineages (Fig 4, Fig S3), which was never observed in the coral host (Davies et al., 2015). Overall our data support the prevailing view that Symbiodiniaceae dispersal is limited, especially relative to their coral hosts, across the seascape. Still, the fact that several MLGs spanned reef sites and even islands highlights the potential for occasional long-range dispersal in *Cladocopium*, especially in C21 (Fig 2E).

## Authors’ Contributions

SWD and MVM conceived of the study, designed the study, coordinated the study and drafted the manuscript. SWD, MRK and MVM collected coral samples. SWD carried out molecular lab work, participated in data analysis, and carried out statistical analyses; DCW, MRK and KM participated in data analysis, statistical analyses and interpretation; All authors gave final approval for publication.

## Acknowledgements

Thanks to field assistants Carly Kenkel, Tim Keitt, Irina Yakushenok and David Stump. Nida Khawaja Rahman contributed to molecular work and James Derry assisted with data management. We acknowledge the Federated States of Micronesia Department of Resources and Development (#11-27-09-01, #11-27-09-02), Guam permit authorities, and the Republic of Palau Bureau of Marine Resources and Koror State Government (Marine Research permit: #09-201; Marine Resource Export Certification #RE-09-23) for all collection permits and CITES export permits. Todd LaJeunesse helped classify *Cladocopium* lineages and provided useful advice on results. We are grateful to Ulrich Mueller, Dan Thornhill, Emily Howells, and the many reviewers who have offered their critical feedback through several sets of revisions throughout the publication process.

## Funding

This study was supported by the Coral Reef Conservation program of the National Oceanic and Atmospheric Administration (NA05OAR4301108) to M.V.M. DCW was funded in part by Pennsylvania State University and grants from the National Science Foundation (OCE-0928764 and IOS-1258058).

## Data Accessibility

All data are available in Supplementary files 1-6 and all data and code used for all analyses and figure generation are publicly available at https://github.com/daviessw/Cladocopium_Micronesia.

## Supplementary Figures

**Fig S1:**
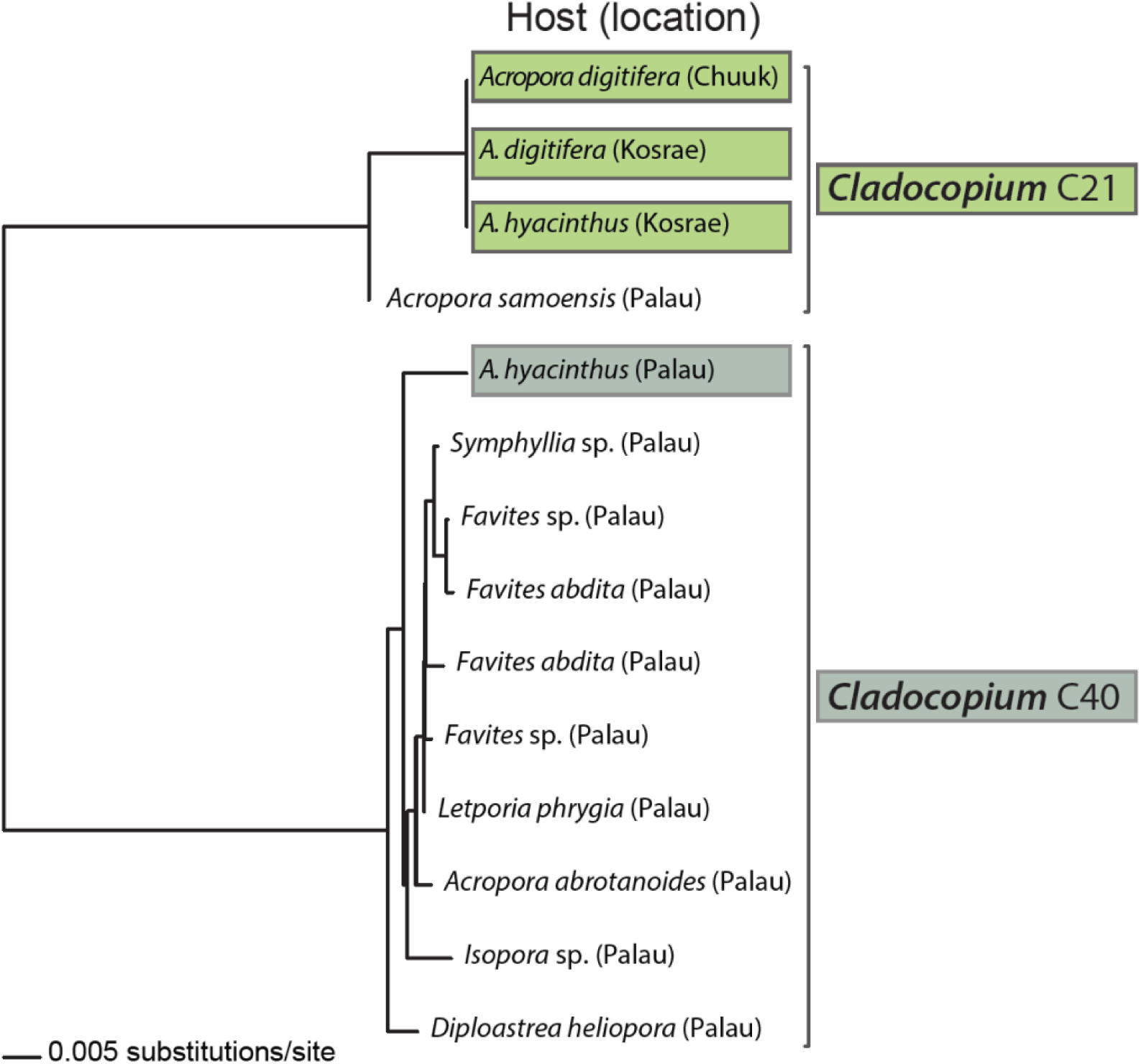
Phylogenetic analysis of psbA_ncr_ sequences for *Cladocopium*. Phylogenetic analysis of psbA_ncr_ sequences from representative samples from this study (indicated by colored boxes) along with publicly available psbA_ncr_ sequences from known *Cladocopium* species identifies the presence of two lineages (C40: Dark green and C21: Light green). The nexus file used to generate the phylogenetic tree is included as Supplemental File S4.

**Fig S2:**
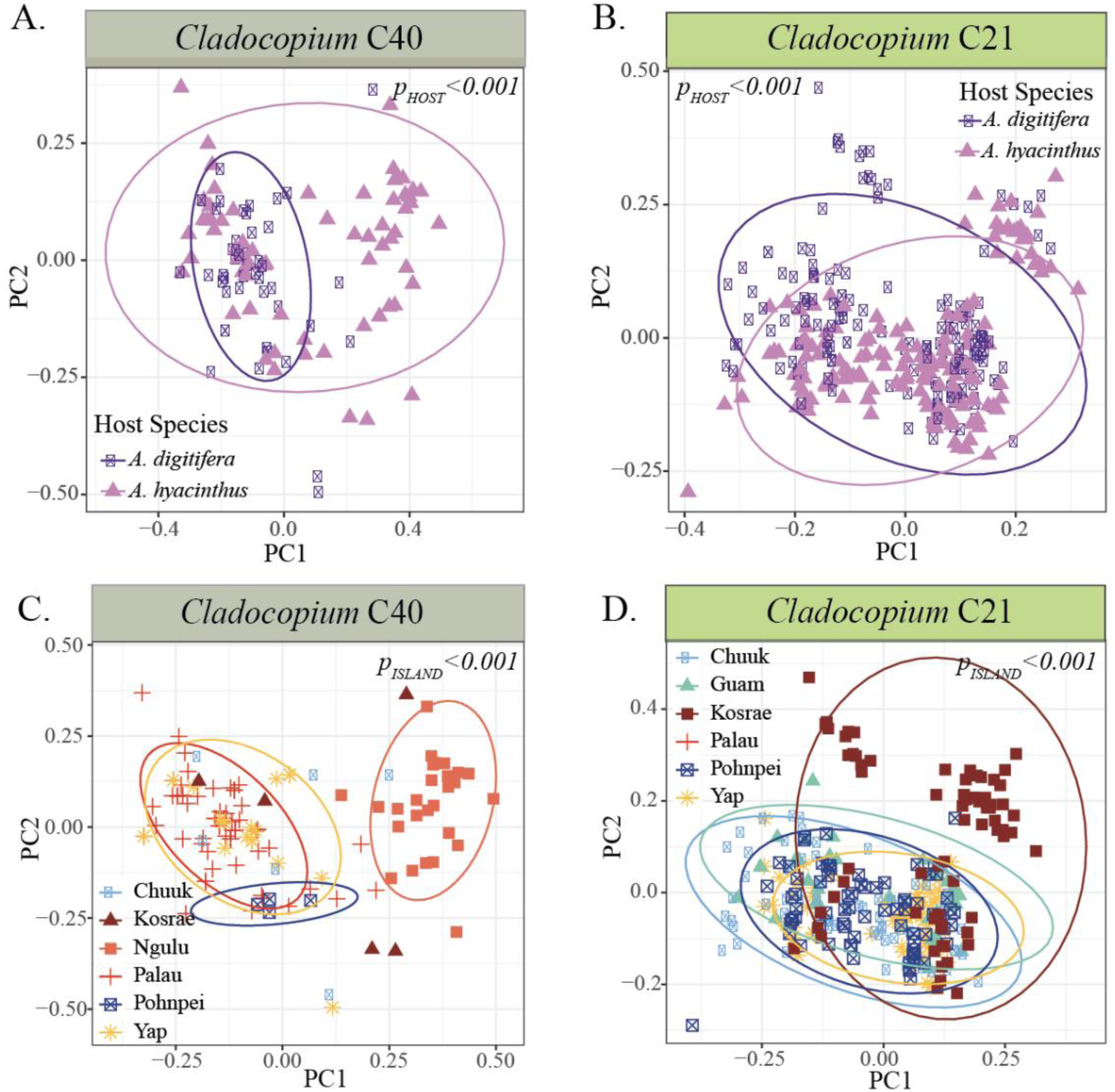
Principle Component Analysis (PCA) of *Cladocopium* binary alleles hosted by *Acropora hyacinthus* and *Acropora digitifera* across seven islands in Micronesia. PCA scatter plots for presence/absence data for all individual samples showing differences between host species in C40 (A), differences between host species in C21 (B), differences between islands in C40 (C), and differences between islands in C21 (D). Sample clustering was significantly different (*adonis*) for host species and islands as indicated by p-values on each panel.

**Fig S3:**
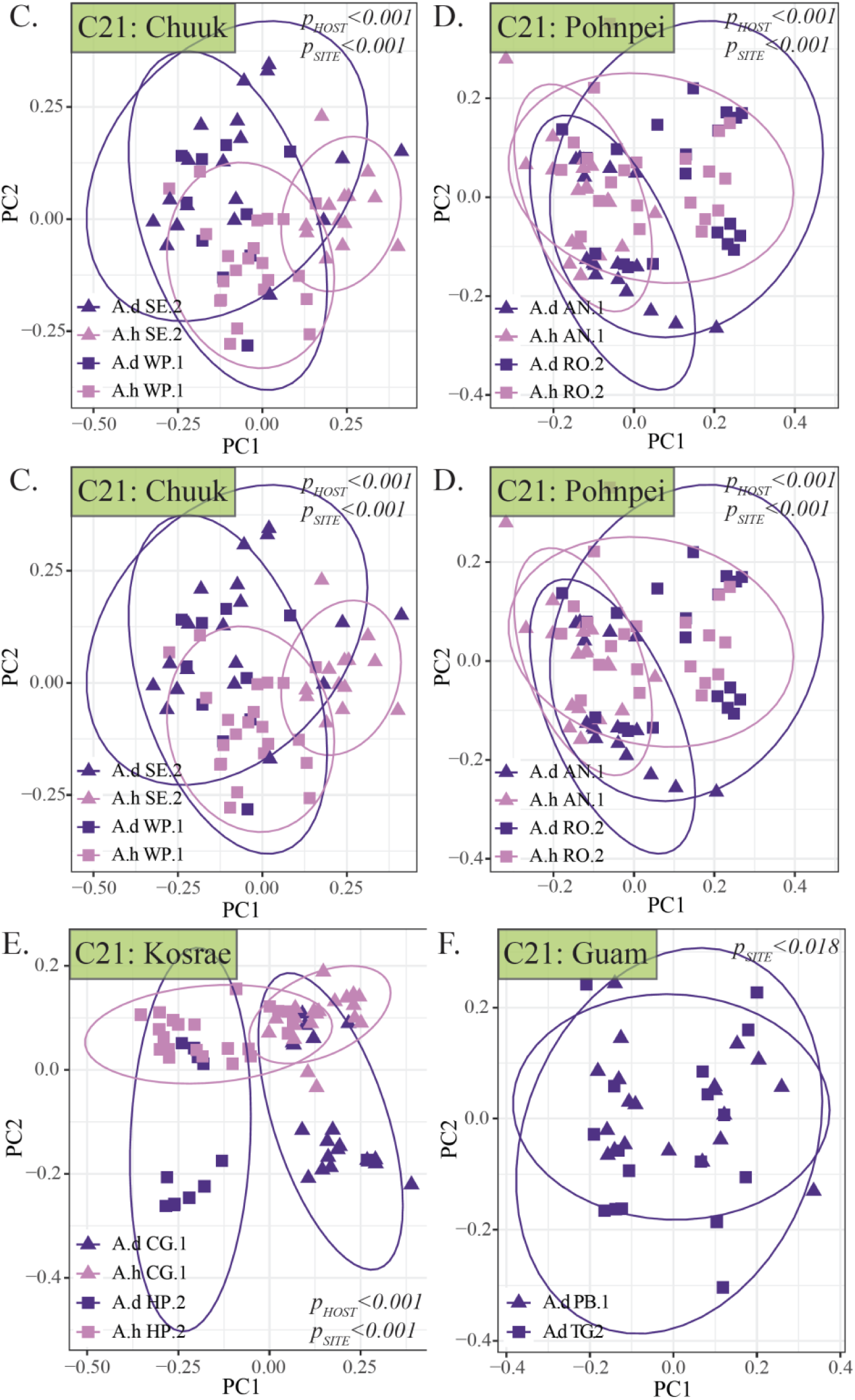
Principle Component Analysis (PCA) of *Cladocopium* binary alleles hosted by *Acropora hyacinthus* and *Acropora digitifera* at twelve reef sites across six islands in Micronesia. PCA scatter plots for presence/absence data for all individual samples showing community clustering within Palau C40 (A), Yap C21 (B), Chuuk C21 (C), Pohnpei C21 (D), Kosrae C21 (E), and Guam C21 (*A. digitifera* hosts only) (F). Sample clustering was significantly different (*adonis*) for host species and site on every island as indicated by p-values on each panel.

## Supplementary Tables

**Table S1:**
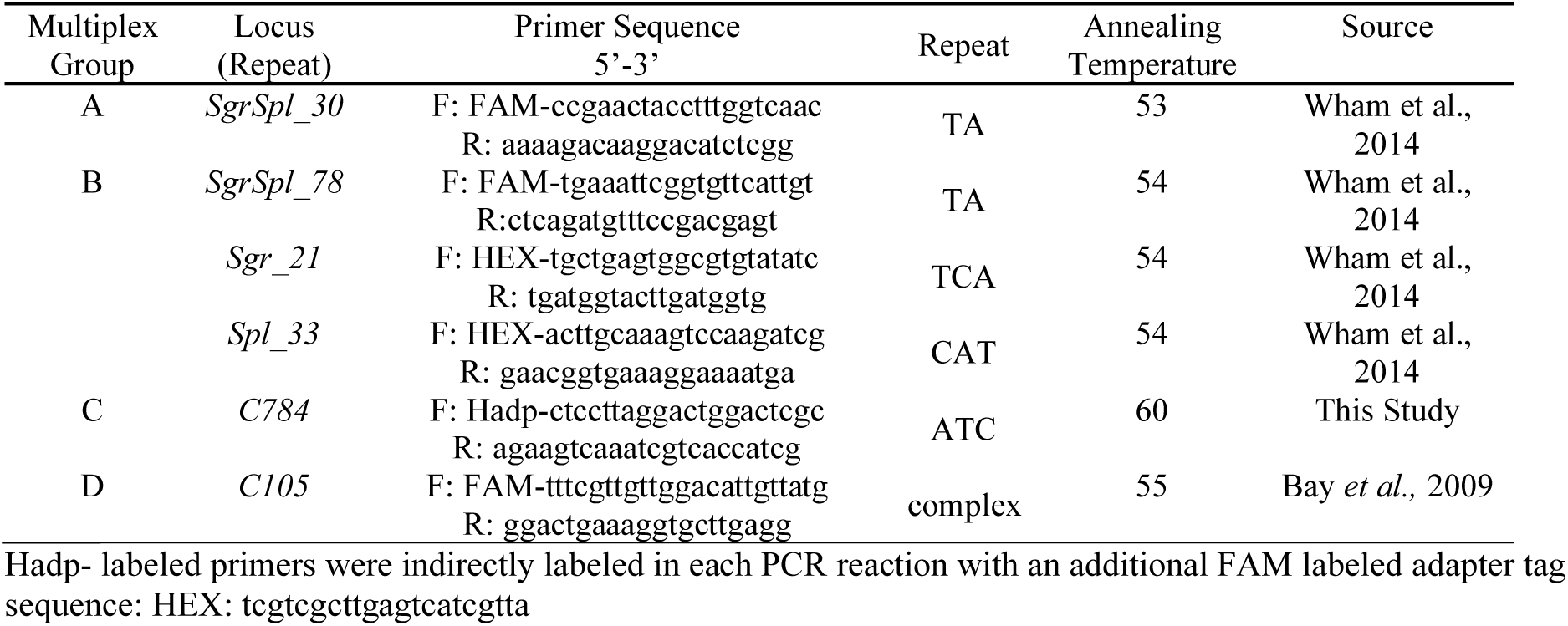
Microsatellite loci summary. Summary of six polymorphic microsatellite loci used to assess genetic variation in *Cladocopium* spp. hosted by *A. hyacinthus* and *A. digitifera* and their corresponding multiplexing groups.

**Table S2:** Discriminant analysis of principle component (DAPC) model information. Discriminant analysis of principle component (DAPC) model information including the number of principle components (“PC”) and discriminant functions (“DF”) retained, the proportion of conserved variance by the clustering model (“var”), and overall assignment proportions across the model (“assign”). A. DAPC information for *Cladocopium* from different host species, B. islands, and C. sites and host species within each island.

**Model Information**
| **A. Host Species** | **PC** | **DF** | **var** | **assign** |
| --- | --- | --- | --- | --- |
| C40 | 16 | 1 | 0.914 | 0.811 |
| C21 | 17 | 1 | 0.915 | 0.838 |
| **B. Islands** | **PC** | **DF** | **var** | **assign** |
| C40 | 17 | 5 | 0.924 | 0.850 |
| C21 | 11 | 5 | 0.804 | 0.704 |
| **C. Within Island** | **PC** | **DF** | **var** | **assign** |
| Palau C40 | 14 | 3 | 0.923 | 0.697 |
| Yap C21 | 16 | 3 | 0.961 | 0.833 |
| Guam C21 | 3 | 3 | 0.522 | 0.810 |
| Chuuk C21 | 10 | 3 | 0.823 | 0.815 |
| Pohnpei C21 | 9 | 3 | 0.814 | 0.846 |
| Kosrae C21 | 7 | 3 | 0.792 | 0.840 |

## Notes

https://github.com/daviessw/Cladocopium_Micronesia

### Competing Interest Statement

The authors have declared no competing interest.

